# Gα13 loss promotes tumor progression in the KPC transgenic mouse model of advanced pancreatic cancer

**DOI:** 10.1101/2021.03.15.435488

**Authors:** Mario A. Shields, Christina Spaulding, Mahmoud G. Khalafalla, Thao N.D. Pham, Hidayatullah G. Munshi

## Abstract

Gα13 transduces signals from G protein-coupled receptors. Gα13 is pro-tumorigenic in epithelial cancer cell lines, which contrasts with its tumor-suppressive function in transgenic mouse models of lymphomas. Here we show that while loss of Gα13 in pancreatic cell lines decreases tumor growth *in vivo*, Gα13 loss in the Kras-driven (KC) mouse model of pancreatic tumor initiation does not affect tumor development or survival. Instead, Gα13 loss in the Kras/Tp53 (KPC) transgenic mouse model of advanced pancreatic cancer promotes well-differentiated tumors with increased tumor burden and reduced survival. Mechanistically, Gα13 loss in the KPC mouse model enhances E-cadherin-mediated cell-cell junctions and mTOR signaling. Importantly, human pancreatic cancers with low Gα13 expression exhibit increased E-cadherin protein expression and mTOR signaling. This work establishes a context-dependent role of Gα13 in pancreatic tumorigenesis, demonstrating a tumor-suppressive role in transgenic mouse models of advanced pancreatic cancer.

## INTRODUCTION

Gα proteins transduce signals from G protein-coupled receptors (GPCRs), the largest class of cell surface proteins that regulate a plethora of biological processes (Oldham and Hamm, 2008). Gα proteins are grouped into Gαs (s, stimulatory), Gαi (i, inhibitory), Gαq, and Gα12 families (Syrovatkina et al., 2016). Mutations in the *GNAS* gene, which encodes Gαs, contribute to the formation of intraductal papillary mucinous neoplasms (IPMNs) (Molin et al., 2013; Wu et al., 2011), precursor lesions of the pancreas that can progress to pancreatic ductal adenocarcinoma (PDAC) tumors (Farrell and Fernandez-del Castillo, 2013; Matthaei et al., 2011). Over 90% of human IPMNs harbor either *KRAS* or *GNAS* mutations or both (Molin et al., 2013; Wu et al., 2011). Co-expression of mutant *Gnas* and mutant *Kras* in mouse models of pancreatic tumorigenesis induces IPMN lesions with a well-differentiated phenotype (Ideno et al., 2018; Patra et al., 2018; Taki et al., 2016). Notably, loss of Tp53 in mouse models of IPMN promotes invasive PDAC lesions (Patra et al., 2018).

While the role of Gαs in pancreatic cancer is now well recognized, the role of other G proteins in pancreatic tumor initiation and progression has yet to be fully defined. Gα13, a member of the Gα12 family, has been implicated in cell migration downstream of GPCRs and receptor tyrosine kinases (Kelly et al., 2007; Kelly et al., 2006; Kozasa et al., 2011) and has been shown to mediate invasion of human PDAC cells (Chow et al., 2016). These results suggest that Gα13 is pro-tumorigenic in many different cancers, including pancreatic cancer. However, studies in B cell lymphomas have found that Gα13 functions as a tumor suppressor (Healy et al., 2016; Muppidi et al., 2014; O’Hayre et al., 2016). The *GNA13* gene is frequently mutated in germinal center-derived B-cell lymphomas, resulting in loss of Gα13 function (Muppidi et al., 2014). Loss of Gα13 in germinal center B cells resulted in the dissemination of these cells and protected them against cell death (Muppidi et al., 2014). Notably, loss of Gα13 in combination with MYC overexpression in germinal center B cells promoted lymphomas (Healy et al., 2016). These results demonstrate that Gα13 functions as a tumor suppressor in lymphomas (Healy et al., 2016; Muppidi et al., 2014; O’Hayre et al., 2016), which contrasts with its function in epithelial cancers (Chow et al., 2016; Kelly et al., 2007; Kelly et al., 2006; Kozasa et al., 2011; Rasheed et al., 2018; Zhang et al., 2018).

It is important to note that many of the studies in epithelial cancers, including our studies in pancreatic cancer (Chow et al., 2016), were conducted in transformed cell lines. However, it is not known whether Gα13 is pro-tumorigenic or tumor-suppressive in transgenic mouse models of epithelial cancer. Here, we evaluate the effects of Gα13 loss in transgenic mouse models of pancreatic tumorigenesis.

## RESULTS

### Gα13 loss does not affect tumor development in the KC transgenic mouse model

To investigate the contribution of Gα13 to epithelial tumor development and progression *in vivo*, we evaluated the effects of Gα13 loss in the LSL-**K**ras^G12D^/Pdx1-**C**re (KC) transgenic mouse model. The KC mouse model utilizes a pancreas-specific Pdx1 promoter-driven Cre recombinase to knock-in the oncogenic mutant Kras^G12D^ into the endogenous locus (Hingorani et al., 2003). Since the Kras mutation is an early event in pancreatic cancer tumorigenesis (Hingorani et al., 2003), the KC mouse model allows us to evaluate the effects of targeting Gα13 on pancreatic tumor initiation (Pham et al., 2021). KC mice were crossed with mice expressing the floxed allele of *Gna13* to generate KCGα13fl/+ and KCGα13fl/fl mice (referred to as KCGfl/+ and KCGfl/fl mice, respectively; **Fig. 1A**). The KCGfl/+ and KCGfl/fl mice showed decreased pancreatic expression of Gα13 at the mRNA and protein levels (**Figs. 1A** and **1B**). There was no effect on *Gna11, Gna12, Gnaq*, or *Gnas* expression in the KCGfl/fl mice compared to KCG+/+ mice (**Fig. 1B**). We next evaluated the effects of Gα13 loss on the lesion burden, proliferation (Ki67), and apoptosis (cleaved caspase 3) in the KCGfl/+ and KCGfl/fl mice. We found no difference in the lesion burden in the KCG+/+, KCGfl/+, and KCGfl/fl mice at either 3- or 6-months of age (**Fig. 1C**). There was also no difference in cleaved caspase 3 (data not shown) or Ki67 staining in the KCGfl/fl mice compared to KCG+/+ mice (**Fig. 1D**). Finally, we evaluated the effects of Gα13 loss on survival in the KC mouse model. There was no difference in the survival of KC, KCGfl/+, and KCGfl/fl mice (**Fig. 1E**).

**Figure 1:**
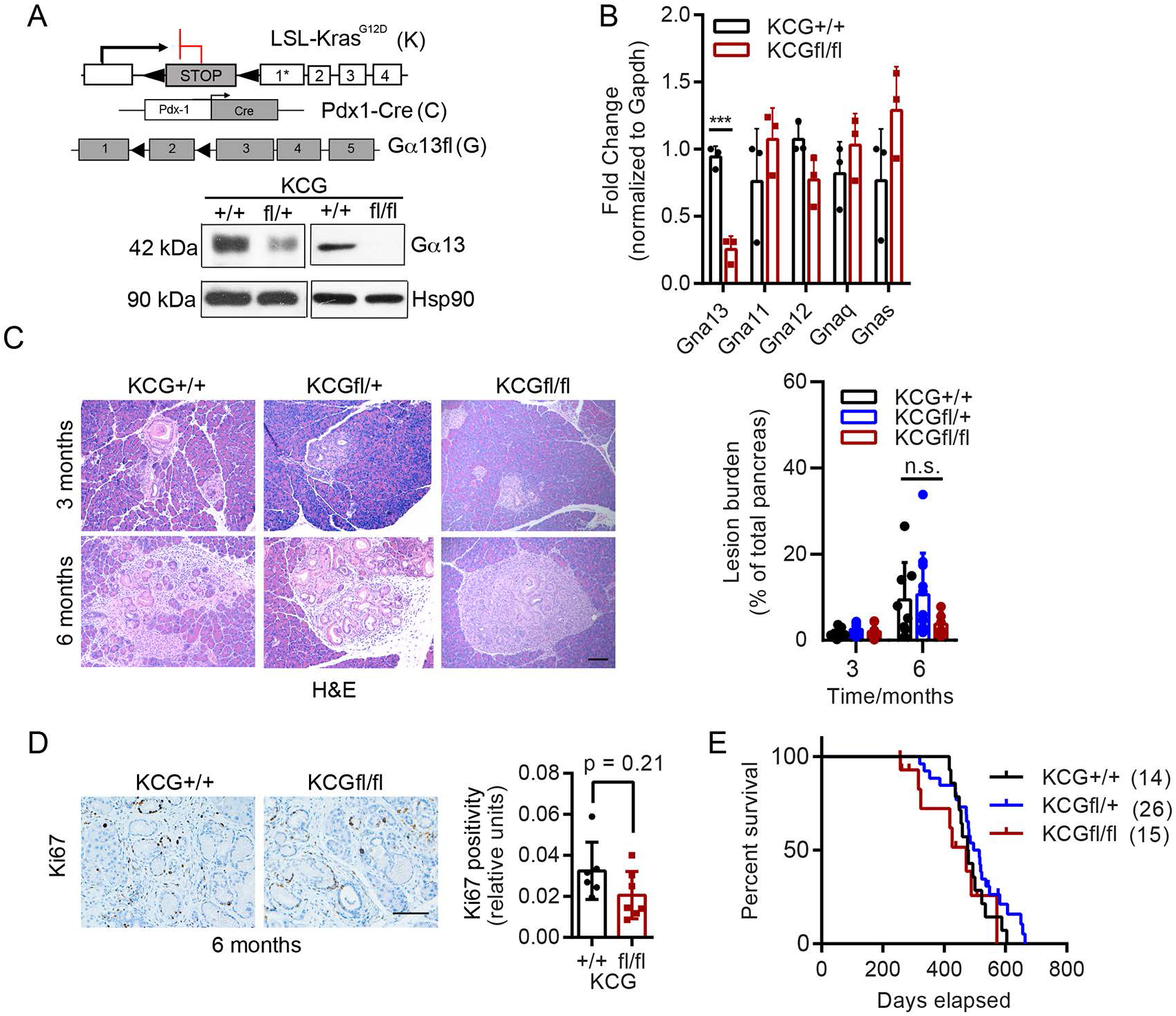
Gα13 loss does not affect tumor development in the KC transgenic mouse model. **A**, Alleles (left) of KrasG12D (K), Pdx1-Cre (C), and Gα13fl (G) and Western blot showing Gα13 expression levels in the pancreas of mice of indicated genotypes. Hsp90 was used as an endogenous control. **B**, Expression of Gna13, Gna11, Gna12, Gnaq, and Gnas was determined by qRT-PCR. Gapdh was used as an endogenous control. t-test (n= 3,3), *** p-value ≤ 0.001, Mean ±SD. **C**, Representative pictures from H&E stains of the pancreas from KCG+/+, KCGfl/+, and KCGfl/fl mice at 3 and 6 months. The whole profile of pancreas tissue was scanned, and the percentage of pancreas with lesions was analyzed at 3 and 6 months (n = 7, 8, 7 at 3 months; n= 8, 11, 6 at 6 months). t-test Mean ±SD. Scale bar = 100 µm. **D**, Immunostains and quantification of Ki67 in the KCG+/+, KCGfl/+, and KCGfl/fl mice at 6 months. t-test, mean ±SD. Scale bar = 100 µm. **E**, Kaplan-Meier survival analysis of KCG+/+ (n=14), KCGfl/+ (n=26), and KCGfl/fl (n=15) mice.

### Cell lines established from KC mice with Gα13 loss grow slower *in vitro* and *in vivo*

Since the previous studies showing Gα13 as pro-tumorigenic were conducted using epithelial cell lines (Chow et al., 2016; Kelly et al., 2007; Kelly et al., 2006; Kozasa et al., 2011; Rasheed et al., 2018; Zhang et al., 2018), we established cell lines from KC, KCGfl/+ and KCGfl/fl mice to evaluate the effects of Gα13 loss. While the KCGfl/+ cell line showed minimal loss of Gα13 protein, the KCGfl/fl cell line showed a near-complete loss of Gα13 (**Fig. 2A**). The KCGfl/fl cell lines showed decreased ability to grow on 2D surfaces (**Fig. 2B**) and reduced ability to grow and invade in 3D culture conditions (**Fig. 2C**). When these cell lines were injected into the flank of B6 mice, the KCGfl/+ cells grew slower than the KCG+/+ cells (**Fig. 2D**). Notably, the KCGfl/fl cells failed to develop tumors *in vivo* (**Fig. 2E**).

**Figure 2:**
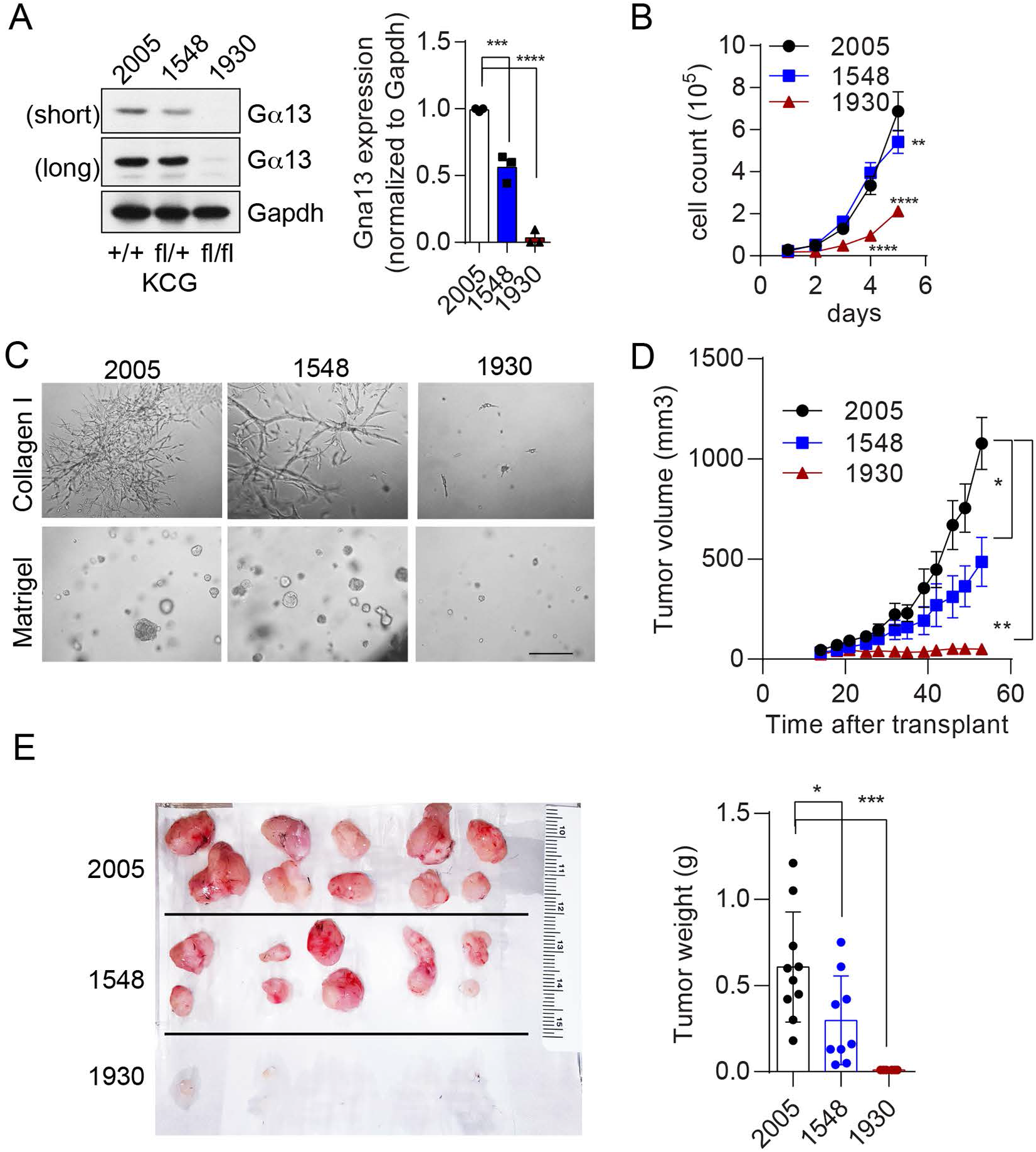
Cell lines established from KC mice with Gα13 loss grow slower *in vitro* and *in vivo*. **A**, Western blot for Gα13 (short and long exposure) in pancreatic cancer cell lines established from KCG+/+ (2005), KCGfl/+ (1548), and KCGfl/fl (1930) mice. **B**, Equal numbers (5 x 10^3^) of KCG+/+ (2005), KCGfl/+ (1548), and KCGfl/fl (1930) cells were plated in 6-well tissue culture plates in duplicate, and proliferation was determined by counting daily the number of cells present. This experiment was repeated at least three times. **C**, Representative picture of KCG+/+ (2005), KCGfl/+ (1548), and KCGfl/fl (1930) cells grown in 3D type I collagen (top) and Matrigel (bottom) cultures after 5 days. Scale bar =100 µm. **D**,**E**, KCG+/+ (2005), KCGfl/+ (1548), and KCGfl/fl (1930) cells were implanted subcutaneously in the flank of female B6 mice (6-8 weeks old, n=5; two tumors per mouse). The tumor sizes were measured using a caliper (***D***), harvested ∼ day 50, photographed, and wet-weight was measured (***E***). One-way ANOVA mean ±SD * p-value ≤ 0.05, ** p-value ≤ 0.01, *** p-value ≤ 0.001; **** p-value ≤ 0.0001.

### Expression of Gα13 shRNA in pancreatic cell lines decreases tumor growth *in vivo*

To further assess the effects of targeting Gα13 in established cell lines, we generated doxycycline-inducible KC cell lines expressing either shRNA against GFP (control) or Gα13 to (**Supplemental Fig. S1A**). When these cell lines were injected in the flank or in the pancreas, loss of Gα13 significantly decreased the growth of KC cell lines *in vivo* (**Supplemental Figs. S1B** and S**1C**).

### KPC mice with Gα13 loss develop differentiated tumors with increased E-cadherin expression

We next evaluated the extent to which Gα13 loss affects tumor progression in the KPC mouse model. The KPC mouse model is a well-established model that expresses mutant Kras and mutant p53 in the mouse pancreas (Hingorani et al., 2005a). We generated KPC mice with heterozygous loss of Gα13 in the mouse pancreas (referred to as KPCGfl/+; **Fig. 3A**). The KPCGfl/+ mice showed decreased pancreatic expression of Gα13 (**Fig. 3B**). At 3 months of age, the KPCGfl/+ developed increased lesion burden compared to the KPCG+/+ mice (**Fig. 3C**). Notably, at the survival endpoint, the KPCGfl/+ mice developed well-differentiated tumors with increased E-cadherin staining and increased proliferation (**Figs. 3D-F**).

**Figure 3:**
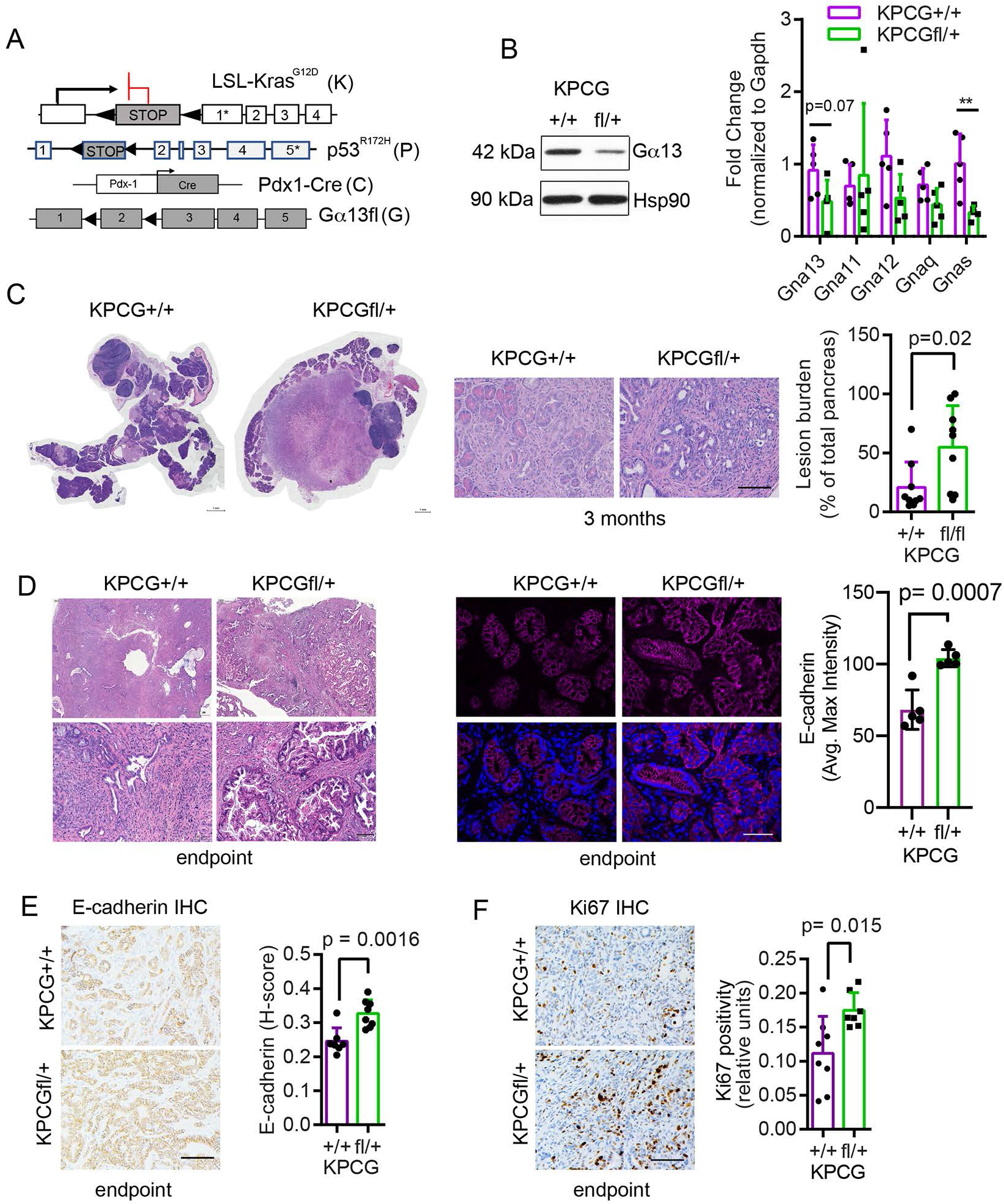
KPC mice with Gα13 loss develop differentiated tumors with increased E-cadherin expression. **A**, Alleles of KrasG12D (K), Pdx1-Cre (C), Gα13fl (G), and Trp53R172H/+ (P). **B**, Gα13 expression levels in the pancreas of mice of indicated genotypes were determined by Western blotting. Hsp90 was used as an endogenous control. Expression of Gna13, Gna11, Gna12, Gnaq, and Gnas was determined by qRT-PCR. Gapdh was used as an endogenous control. Unpaired t-test (n= 3,3), ** p-value ≤ 0.01. Mean ±SD. **C**, Representative pictures (left) from H&E stains of the pancreas from KPCG+/+ and KPCGfl/+ mice at 3 months. The whole profile of pancreas tissue was scanned, and the relative lesion burden and the lesion type were quantified at 3 months (n = 9, 9). t-test Mean ±SD. Scale bar = 100 µm. **D**, H&E and immunofluorescence stains for E-cadherin in the pancreatic tissue from KPCG+/+ and KPCGfl/+ mice at the endpoint. The relative fluorescence intensity at cell-cell junctions in the lesions was quantified using ImageJ. **E**, Immunohistochemical stains for E-cadherin in the pancreatic tissue from KPCG+/+ and KPCGfl/+ mice at the endpoint. The relative E-cadherin score at cell-cell junctions in the lesions was quantified using HistoQuest Software. **F**, Immunostains and quantification of Ki67 in the KPCG+/+ and KPCGfl/+ mice (n= 8, 7). t-test, mean ±SD. Scale bar = 100 µm.

### KPCGfl/+ mice demonstrate increased mTOR signaling and decreased survival

To understand why the KPCGfl/+ tumors show increased proliferation, we analyzed human data to identify signaling pathways in human PDAC tumors associated with decreased Gα13 expression. We initially evaluated the relationship between E-cadherin and Gα13 in human PDAC tumors using the publicly accessible The Cancer Genome Atlas (TCGA) data in the cBioportal (**Supplemental Table S1**) (Cerami et al., 2012; Gao et al., 2013). Human PDAC tumors with low Gα13 expression showed increased E-cadherin protein expression, but there was no difference in *E-cadherin* mRNA levels (**Fig. 4A**). There was increased expression of β-catenin (CTNNB1) and claudin-7 (CLDN7) in human tumors with reduced Gα13 expression (**Fig. 4A**). There was also evidence of increased activation of mTOR signaling, with increased protein expression of RPS6, p-PDK1(S241), EEF2, and MTOR in human PDAC tumors with reduced Gα13 expression (**Fig. 4B**). Human PDAC tumors with reduced Gα13 expression showed no difference in *RPS6* mRNA levels, but a statistically significant reduction in *MTOR* mRNA levels (**Fig. 4B**). Notably, while we did not see increased levels of p-PDK1 in tumors developing in the KPCGfl/+ tumors compared to KPC tumors (data not shown), we found that there was increased expression of Rps6 and Mtor in the KPCGfl/+ tumors compared to KPCG+/+ mice (**Fig. 4C**). Finally, we found significantly reduced survival of KPCGfl/+ mice compared to KPC mice (**Fig. 4D**). These results suggest a tumor-suppressive role for Gα13 in advanced PDAC tumors.

**Figure 4:**
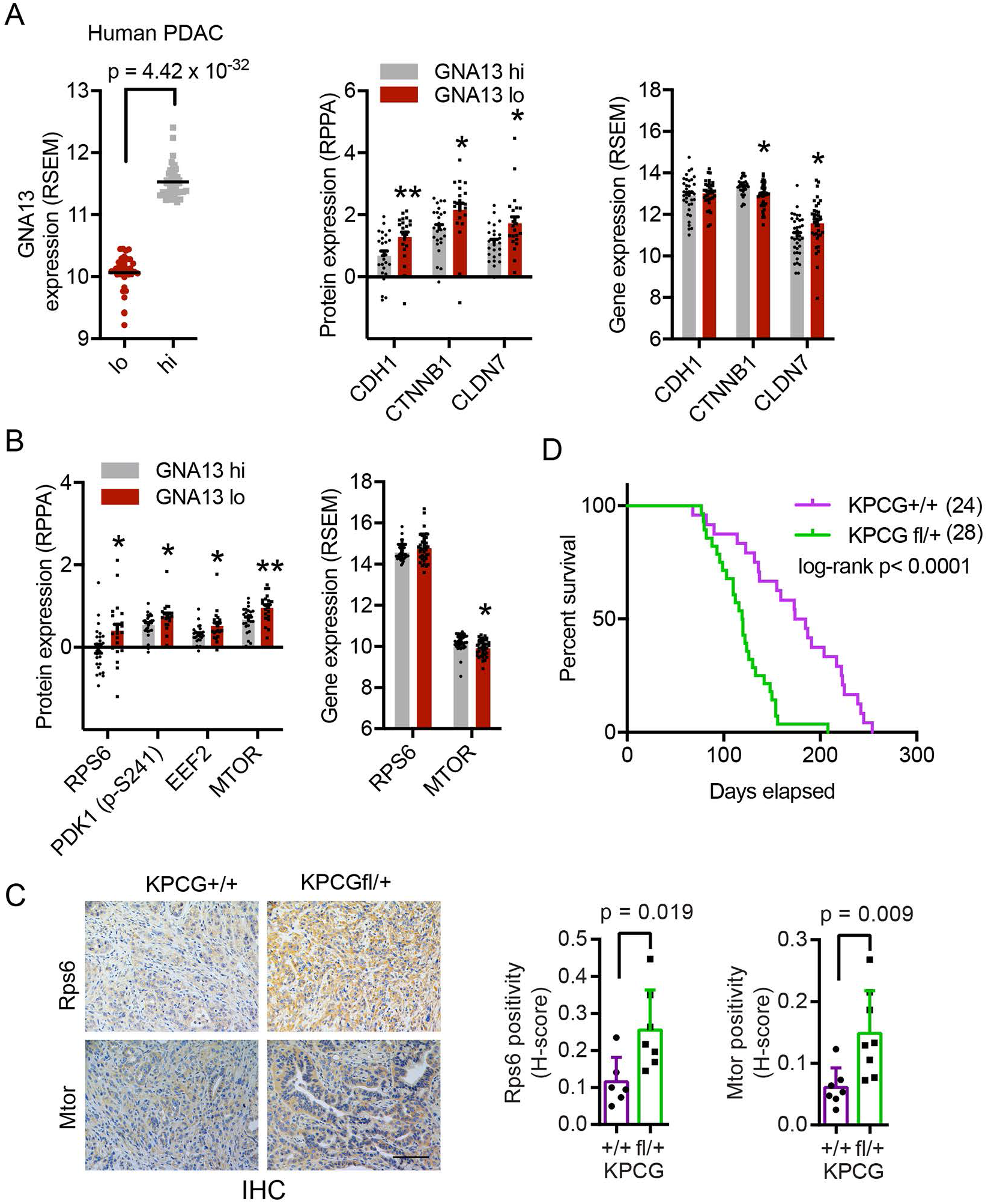
KPCGfl/+ mice demonstrate increased mTOR signaling and decreased survival. **A**, Analysis of samples in cBioportal with low and high expression of *GNA13* for E-cadherin (CDH1), β-catenin (CTNNB1), and Claudin-7 (CLDN7) expression at the protein (RPPA) and gene expression (RSEM) levels (n= 43, 43). t-test, mean ±SEM. **B**, Analysis of samples in cBioportal with low and high expression of *GNA13* for RPS6, PDK1 (p-S241), EEF2, and MTOR at the protein (RPPA) levels. Effects on RPS6 and MTOR were also analyzed at the gene expression (RSEM) levels. t-test, mean ±SEM *, p-value ≤ 0.05; **, p-value ≤ 0.01. **C**, Immunostains and quantification for Rps6 (n=6,6) and Mtor (n=7,7) in advanced tumors developing in KPCG+/+ and KPCGfl/+ mice. t-test, mean ±SD. Scale bar = 100 µm. **D**, Kaplan-Meier survival analysis of KPCG+/+ (n=21) and KPCGfl/+ (n=28) mice using long-rank test.

## DISCUSSION

GPCRs are targets of several currently approved therapies (Hauser et al., 2017). There is increasing interest in understanding the role of effectors downstream of GPCRs and determining whether these effectors could be potential therapeutic targets (Hauser et al., 2017). Since GPCRs can signal through the G12 family of small G proteins (Kurose, 2003), many studies have evaluated the contribution of Gα12 and Gα13 to physiologic and pathologic processes (Sriram et al., 2020; Syrovatkina and Huang, 2019; Tutunea-Fatan et al., 2020). These studies, which have been primarily performed in epithelial cancer cell lines, showed that Gα13 functions as a tumor promoter (Chow et al., 2016; Kelly et al., 2007; Kelly et al., 2006; Kozasa et al., 2011; Rasheed et al., 2018; Zhang et al., 2018). We also now show that targeting Gα13 in Kras-expressing pancreatic cell lines reduces or blocks tumor growth *in vivo*. However, targeting Gα13 in the Kras-driven KC transgenic mouse model of tumor initiation did not affect tumor development or survival of these mice. Instead, we have found that targeting Gα13 in the Kras and p53-driven KPC mouse model promoted tumor development and decreased survival, indicating that Gα13 has a tumor-suppressive role in advanced tumors.

Previously published studies in B cell lymphomas have also shown that Gα13 functions as a tumor suppressor (Healy et al., 2016; Muppidi et al., 2014; O’Hayre et al., 2016). The *GNA13* gene is frequently mutated in germinal center-derived B-cell lymphomas, resulting in loss of Gα13 function (Muppidi et al., 2014). Loss of Gα13 in germinal center B cells resulted in the dissemination of these cells and protected them against cell death (Muppidi et al., 2014). Loss of Gα13 in combination with MYC overexpression in germinal center B cells promoted lymphomas (Healy et al., 2016). These results in a lymphoma transgenic mouse and our data in the KPC transgenic mouse model demonstrate a tumor-suppressive role of Gα13 in tumor development and progression.

Notably, Gα13 loss parallels many of the effects of gain-of-function mutations in Gnas in the pancreas. Co-expression of mutant Gnas with mutant Kras promoted the development of IPMNs (Ideno et al., 2018; Patra et al., 2018; Taki et al., 2016). However, the expression of mutant Gnas in Kras-expressing cell lines decreased the number of colonies in soft agar assay, reduced invasion *in vitro*, reduced tumor growth *in vivo*, and induced striking epithelial differentiation (Ideno et al., 2018). Similar to the expression of mutant Gnas in cell lines, we have found that loss of Gα13 in Kras-expressing cell lines decreased proliferation *in vitro*, decreased growth and invasion in 3D collagen and matrigel, and decreased tumor growth *in vivo*. In contrast, while co-expression of mutant Kras and Gnas was sufficient to induce IPMNs (Ideno et al., 2018; Patra et al., 2018; Taki et al., 2016), we did not see any apparent effects of Gα13 loss on IPMN development in the KC mouse model. However, when we targeted Gα13 in the KPC mouse model, we found that loss of Gα13 resulted in differentiated tumors, similar to the phenotype seen with co-expression of Gnas and Kras (Ideno et al., 2018; Patra et al., 2018; Taki et al., 2016). We found that mice with Gα13 loss in the KPC model died faster than the control KPC mice, likely from the extensive tumor burden despite the tumors being well-differentiated. Our findings are similar to the findings in the Kras/Gnas model, where the mice also died faster (Ideno et al., 2018; Taki et al., 2016).

We show increased E-cadherin expression at cell-cell junctions in the KPC mouse model, thus explaining the well-differentiated tumors in the KPC mice with Gα13 loss. We had previously shown that Gα13 loss results in increased E-cadherin at cell-cell junctions (Chow et al., 2016). Loss of Gα13 has also been shown to decrease endocytosis of VE-cadherin, resulting in increased VE-cadherin at cell-cell junctions (Gong et al., 2014). Earlier studies have shown that E-cadherin undergoes endocytosis following the induction of pancreatitis and subsequent recycling to regenerate cell-cell junctions and mediate acinar cell recovery (Lerch et al., 1997). Importantly, human PDAC tumors with Gα13 loss also demonstrated increased expression of E-cadherin and other cell adhesion proteins. Human PDAC tumors with decreased expression of Gα13 also exhibited increased activation of the mTOR signaling pathway. In agreement with the human data, we also found that Gα13 loss in the KPC mouse model enhanced mTOR signaling. Notably, GPCRs and G proteins have been shown to modulate mTOR signaling (Gan et al., 2019; Jewell et al., 2019; Nagai et al., 2020; Puertollano, 2019). For example, Gαs-coupled GPCRs activate protein kinase A to inhibit mTOR complex 1 in multiple cell lines and mouse tissue (Jewell et al., 2019).

Overall, our findings increase our understanding of the role of Gα13 *in vivo*. We show that the phenotypic changes seen with Gα13 loss parallel many of the phenotypic changes seen with the expression of mutant Gnas (Ideno et al., 2018; Patra et al., 2018; Taki et al., 2016). However, in contrast to the previously published findings that Gα13 functions as a tumor promoter in established cell lines (Chow et al., 2016; Kelly et al., 2007; Kelly et al., 2006; Kozasa et al., 2011; Rasheed et al., 2018; Zhang et al., 2018), we demonstrate that Gα13 has a tumor-suppressive role in advanced PDAC tumors in transgenic mouse models.

## METHODS

### Animal experiments

#### Study approval

All animal work and procedures were approved by the Northwestern University Institutional Animal Care and Use Committee. In addition, all animal experiments were performed in accordance with relevant guidelines and regulations.

#### Conditional knockout

Mice with loss of Gα13 in the pancreas were generated by crossing Pdx1-Cre mice (Jackson Laboratory #014647) to mice expressing the floxed allele of *Gna13* (kindly provided by Stefan Offermans, Max Planck Institute) (Moers et al., 2003), to generate CGα13fl/+ and CGα13fl/fl mice. The bigenic mice were further crossed with mice expressing an LSL-KRas^G12D^ (Jackson Laboratory #019104) mutant allele to generate KCGα13fl/+ (KCGfl/+) or KCGα13fl/fl (KCGfl/fl) mice. These mice were further crossed with mice expressing LSL-Trp53^R172H/+^ (Jackson Laboratory 008652) to generate KPCGfl/+ mice. All mice were bred on a C57/BL6 background.

#### Cancer cell implantation

For orthotopic implantation of cancer cells, a small (∼1 cm) left abdominal side incision through skin and peritoneum of an anesthetized C57BL/6 mouse was made, and cells (1 × 10^5^) suspended in 25 µL of growth factor-reduced Matrigel (Corning Inc., Corning, NY) were injected into the tail of the pancreas using a 22-gauge Hamilton needle. The incision in the peritoneum was closed with absorbable VICRYL polyglactin sutures (Ethicon, Somerville, NJ), and the skin was closed with 7 mm Reflex wound clips (Roboz, Gaithersburg, MD). Tumor burden in mice was measured weekly using bioluminescence imaging using the Lago Bioluminescence system (Spectral Instrument Imaging, Tucson AZ). Mice were intraperitoneally administered D-luciferin potassium salt (GoldBio, St. Louis, MO) at a dose of 150 mg/kg, and the bioluminescence signal was acquired within 10-15 minutes after injection.

For subcutaneous implantation, cancer cells (1 × 10^5^) were suspended in 100 µL of 1:1 mixture of PBS: Matrigel was injected under the skin in the flank of C57BL/6 mice. For shRNA induction, mice were fed water containing doxycycline (200 mg/L), which was changed every 2-3 days. Tumor dimensions were measured using a digital caliper, and volume was determined using the formula V = (L × W^2^)/2, where V is the volume, L is length, and W is the width.

#### Endpoint

At the experimental endpoint, mice were euthanized, pancreatic tissue weighed, and then fixed in 10% neutral buffered formalin overnight or flashed frozen for RNA or protein extraction. Fixed tissue was subsequently processed and embedded in paraffin for histological stains and immunohistochemistry.

### Murine pancreatic cell lines

The mouse pancreatic cell line KC4868, derived from genetically engineered *Kras*^*LSL-G12D/+*^; *Pdx1-Cre* (KC) C57BL/6 mice (Hingorani et al., 2005b), was provided by Dr. Paul Grippo (University of Illinois, Chicago, USA). The KC4868 cells were determined to be free of mycoplasma using MycoAlert Plus kit (Lonza Walkersville, MD) and routinely maintained in DMEM with 10% fetal bovine serum (FBS), 1% penicillin/streptomycin.

#### Cell lines expressing inducible Gα13 shRNA

KC4868 cells expressing doxycycline-inducible short hairpin RNA (shRNA) against Gα13 were generated using a lentiviral vector-based method. Four hairpins sequences against *Gna13* were obtained from the RNAi Consortium database and purchased from Integrated DNA Technologies (Coralville, IA). The most potent shRNAs were found to be TRCN0000098145 and TRCN00000435189, and these were used along with control shRNA against eGFP (5’-CCGGTACAACAGCCACAACGTCTATCTCGAG-ATAGACGTTGTGGCTGTT GTATTTTTG-3’) for subsequent experiments. Cancer cells expressing doxycycline-regulated shRNAs against *Gna13* were generated using the Tet-pLKO-puro lentiviral system, deposited by Dr. Dmitri Wiederschain to Addgene (Addgene plasmid #21915), by following their protocol (Wiederschain et al., 2009). Lentiviral particles were generated in 293T cells, and puromycin (2 µg/mL) was used to select transduced cells. Gene knockdown was induced by incubating cells with 1 µg/mL of doxycycline for 24 to 48 hours. Expression of luciferase in cancer cells was achieved by transducing cells with retroviral particles, generated in Phoenix-Eco cells, then selected using blasticidin (10 µg/mL).

#### Generation of cell lines from transgenic mouse model

KC, KCGfl/+ or KCGfl/fl mice more than 6 months of age were euthanized and the harvested pancreatic tissue placed in 5 ml of cold DMEM using aseptic technique. The tissue was minced using a sterile scalpel and then placed in a 2 mg/ml Collagenase I solution diluted in DMEM and enzymatically digested at 37^°^C for 30 minutes with mixing by vortex at 10-minute intervals. The digested tissue was centrifuged at 1200 rpm for 3 minutes and the cell pellet resuspended in DMEM media with 10% FBS and 1% penicillin/streptomycin. The cells were plated in a 10-cm tissue culture dish and maintained at 37^°^C with 5% CO_2_ in an incubator. The cells were serially passaged until the culture was purely cancer cells (at passage 4 or 5). For 3D matrix growth, cancer cells (5 × 10^3^) were suspended in 500 µl of Matrigel (cat # 354234 Corning) or neutralized rat tail collagen type I (cat # 354236 Corning) at 2 mg/ml.

### Immunohistochemistry

For immunostains, paraffin-embedded sections were deparaffinized and rehydrated. Antigen retrieval was performed by boiling for ten minutes in sodium citrate buffer (10 mM, pH 6.0) using a pressure cooker. Endogenous peroxidase activity in tissue was quenched, and sections were blocked with a mixture of goat serum and bovine serum albumin (BSA). Tissue sections were incubated with antibodies: for Ki-67 (Cell Signaling #12202, 1:500), cleaved caspase-3 (Cell Signaling #9664, 1:800), MTOR (Cell Signaling #2983, 1:500), and RPS6 (Cell Signaling #2217, 1:1000) overnight at 4^°^C. Antibody binding was detected using HRP-conjugated anti-rabbit secondary antibody and visualized using ImmPACT DAB Peroxidase Substrate kit (VectorLabs, Burlingame, CA). Photographs were taken on the FeinOptic microscope with a Jenoptik ProgRes C5 camera or TissueGnostics system and analyzed by ImageJ or HistoQuest Software.

### Immunofluorescence

After antigen retrieval (similar to IHC), tissue sections were stained using the mouse-on-mouse (MOM) immunodetection kit (Vector Laboratories (BMK-2202). Tissue sections were blocked with MOM-blocking reagent for 1 hour at room temperature, followed by washing and incubation with MOM-diluent for 5 minutes. Tissue sections were then incubated with primary a primary anti-E-cadherin (BD Biosciences #610181, 1:200) antibody for 1 hour. After washing, the sections were incubated with Alexa Fluor conjugated goat-anti mouse (AF647, #A21235) secondary antibody for 30 minutes in the dark at room temperature. The tissue sections were washed and incubated with DAPI (Thermo Fisher D1306) for 10 minutes. Slides were mounted with fluorescence mounting media and images acquired using the EVOS M5000 microscope (Thermo Fisher).

### Western blot

Tissue samples were finely ground using a cold mortar and pestle. Protein was isolated from cultured cells or ground tissue using ice-cold RIPA lysis buffer containing protease and phosphatase inhibitors. The lysates were then clarified by centrifugation at 10,000 rpm for 10 minutes at 4^°^C, and the protein concentration was determined using Precision Red solution (Cytoskeleton, Inc., Denver, CO) according to the manufacturer’s instructions. Equal amounts of protein were separated with a 10% SDS-PAGE electrophoresis gel. The separated proteins were transferred to a nitrocellulose membrane using the semi-dry transfer system (Bio-Rad). After blocking for one hour at room temperature with 5% BSA, the membranes were incubated overnight at 4^°^C with primary antibodies. Primary antibodies used include Gα13 (Santa Cruz sc-293424), E-cadherin (Cell Signaling 3195), Gapdh (Millipore Sigma #MAB374), Hsp90 (Santa Cruz Biotechnology sc-7947), and alpha-tubulin (Santa Cruz Biotechnology sc-8035). HRP-conjugated rabbit (A6667) or mouse (A4416) secondary antibody (Millipore-Sigma St. Louis, MO) was used with SuperSignal West Pico PLUS (Thermo Fisher Scientific) for protein detection.

### qRT-PCR

RNA was isolated from cultured cells or tissue using the RNeasy kit (74104, QIAGEN, Hilden, Germany) according to the manufacturer’s instructions. cDNA was synthesized from 1-2 µg of RNA with random hexamers and M-MLV reverse transcriptase in a single 50 µL reaction. Quantitative PCR was performed using gene-specific TaqMan probes, TaqMan Universal PCR Master Mix, and the CFX Connect Real-Time PCR Detection System (Bio-Rad Hercules, CA). Probes used to detect the following genes: *Gna13* (Mm01250415_m1), *Gna11* (Mm01172792_m1), *Gna12* (Mm00494665_m1), *Gnaq* (Mm00492381_m1), *Gnas* (Mm01242435_m1). Mouse *Gapdh* (Mm99999915_g1) and *Rn18s (*Mm03928990_g1) were used as endogenous controls, and their expression levels were used to calculate the relative expression using the comparative C_t_ method (Livak and Schmittgen, 2001).

### TCGA data analysis

Expression data from RNA and protein (RPPA) analyses for human pancreatic ductal adenocarcinoma patients in The Cancer Genome Atlas (TCGA) were extracted from cBioportal (Cerami et al., 2012; Gao et al., 2013). Patient IDs from the study are listed in Table 1 and are categorized as high (GNA13 hi, upper 25%, n= 43) or low (GNA13 lo, lower 25%, n = 43) expression of *GNA13* RNA.

### Statistics

The *in vivo* and *in vitro* results were compared using one-way ANOVA and 2-tailed *t*-test analysis. Error bars represent the standard error of the mean or standard deviation as specified in the figure legends. All statistical analyses were done using GraphPad Prism. A p-value of less than 0.05 was considered significant.

## ACKNOWLEDGEMENTS

We thank Dr. Stefan Offermanns (Max-Planck-Institute for Heart and Lung Research, Bad Nauheim, Germany), who kindly provided the *Gna13*^*flox/flox*^ mice. This work was supported by grants R01CA217907 (to H.G.M.), R21CA255291 (to H.G.M.), a Merit award I01BX002922 (to H.G.M.) from the Department of Veterans Affairs, APA/APA Foundation 2020 Young Investigator Pancreatitis Grant (to M.A.S.), the Mander Foundation Award (to H.G.M.) and the Harold E. Eisenberg Foundation Award (to T.N.D.P.) from the Robert H. Lurie Cancer Center, and the NIH/NCI training grant T32 CA070085 (to T.N.D.P).

## AUTHOR CONTRIBUTIONS

M.A.S. designed the studies, performed the experiments, analyzed the data, and wrote the manuscript. C.S., M.G.K., and T.N.D.P. performed the experiments and analyzed the data. H.G.M. designed the studies, analyzed the data, wrote the manuscript, and secured funding. All authors edited and approved the final manuscript.

## SUPPLEMENTAL INFORMATION

**Supplemental Figure S1: G**α**13 shRNA in KC cell lines decreases tumor growth *in vivo*. A**, 4868 KC cells expressing doxycycline-inducible short hairpin RNA against GFP or Gα13 were treated with doxycycline (1 µg/ml) and analyzed by western blot (top, 48h) and qRT-PCR (bottom, 24h). α-tubulin and Gapdh were used as an endogenous control for western blot and qRT-PCR, respectively. The experiment was conducted at least three times. t-test, Mean ±SD. **, p<0.01; ***, p<0.001. **B**, 4868 KC cells (1 × 10^4^) expressing shGFP or shRNA against Gα13 (sh45 or sh89) were implanted subcutaneously in the flank of female mice (6-8 weeks old). After 2 weeks, mice were fed doxycycline containing water (200 mg/L). The tumor sizes were measured using a caliper until an endpoint was reached, then harvested and stained using H&E. One-way ANOVA (n=3,3,2; two tumors per mouse). Mean ±SD. Scale bar = 1mm. **C**, 4868 KC cells (1 × 10^4^) expressing shGFP or shRNA against Gα13 (sh45) and co-expressing luciferase were implanted in the pancreas of mice at 8 weeks of age. Tumor burden was measured weekly using bioluminescence imaging. Shown here are images after 35 days. At the endpoint, tumors were harvested, wet-weight measured, and then stained using H&E. (n=3, 3) t-test Mean ±SD. Scale bar = 1mm.

**SUPPLEMENTAL TABLE S1:** Expression data from RNA and protein (RPPA) analyses for human pancreatic ductal adenocarcinoma patients in The Cancer Genome Atlas (TCGA) were extracted from cBioportal. Expression of *GNA13* RNA is categorized as high (GNA13 hi, upper 25%, n= 43) or low (GNA13 lo, lower 25%, n = 43). Table 1 lists patient IDs from TCGA used in our analysis.

