## SupplementalTable1 for "Gα13 loss promotes tumor progression in the KPC transgenic mouse model of advanced pancreatic cancer"

Table 1

| Patient | Expression | Group |
| --- | --- | --- |
| TCGA-3A-A9IJ | 595.29 | Low |
| TCGA-US-A779 | 679.62 | Low |
| TCGA-OE-A75W | 685.2 | Low |
| TCGA-2J-AABV | 811.02 | Low |
| TCGA-3A-A9IR | 870.31 | Low |
| TCGA-FB-A5VM | 871.15 | Low |
| TCGA-FB-AAPP | 907.67 | Low |
| TCGA-FB-AAQ6 | 1018.78 | Low |
| TCGA-YY-A8LH | 1049.06 | Low |
| TCGA-3A-A9I9 | 1050.42 | Low |
| TCGA-IB-A5SP | 1054.38 | Low |
| TCGA-2J-AABE | 1061.82 | Low |
| TCGA-HZ-A77O | 1063.4 | Low |
| TCGA-HV-A5A3 | 1072.22 | Low |
| TCGA-3A-A9IO | 1080.56 | Low |
| TCGA-US-A77G | 1086.72 | Low |
| TCGA-HZ-7289 | 1088.34 | Low |
| TCGA-2L-AAQM | 1093.18 | Low |
| TCGA-F2-6880 | 1115.68 | Low |
| TCGA-3A-A9IN | 1117.21 | Low |
| TCGA-3A-A9IL | 1121.18 | Low |
| TCGA-US-A776 | 1139.15 | Low |
| TCGA-3A-A9IH | 1144.9 | Low |
| TCGA-YH-A8SY | 1168.81 | Low |
| TCGA-3A-A9IS | 1183.55 | Low |
| TCGA-LB-A9Q5 | 1194.79 | Low |
| TCGA-2L-AAQL | 1201.19 | Low |
| TCGA-HZ-8001 | 1245.81 | Low |
| TCGA-F2-A8YN | 1249.43 | Low |
| TCGA-LB-A8F3 | 1254.82 | Low |
| TCGA-FB-AAQ0 | 1280.08 | Low |
| TCGA-IB-AAUM | 1370.61 | Low |
| TCGA-IB-A6UF | 1388.65 | Low |
| TCGA-IB-A6UG | 1394.58 | Low |
| TCGA-2J-AABI | 1396.23 | Low |
| TCGA-IB-7887 | 1407.02 | Low |
| TCGA-HV-A7OL | 1413.43 | Low |
| TCGA-FB-AAQ3 | 1414.74 | Low |
| TCGA-H6-A45N | 1416.21 | Low |
| TCGA-3A-A9IV | 1423.88 | Low |
| TCGA-LB-A7SX | 1427.81 | Low |
| TCGA-RB-AA9M | 1429.61 | Low |
| TCGA-2L-AAQE | 1433.27 | Low |
| TCGA-2J-AABT | 2262.83 | High |
| TCGA-IB-7652 | 2263.45 | High |
| TCGA-3A-A9I5 | 2288.12 | High |
| TCGA-IB-A7LX | 2292.37 | High |

| | | |
|---|---|---|
| TCGA-FB-A7DR | 2298.03 | High |
| TCGA-HZ-7920 | 2328.29 | High |
| TCGA-IB-7885 | 2335.15 | High |
| TCGA-HZ-7924 | 2347.63 | High |
| TCGA-IB-AAUV | 2353.62 | High |
| TCGA-IB-7649 | 2413.89 | High |
| TCGA-YB-A89D | 2421.38 | High |
| TCGA-HZ-A77Q | 2429.49 | High |
| TCGA-3A-A9IZ | 2477.32 | High |
| TCGA-IB-7891 | 2502.84 | High |
| TCGA-FB-AAPS | 2536.55 | High |
| TCGA-FB-A545 | 2541.26 | High |
| TCGA-IB-AAUR | 2591.52 | High |
| TCGA-IB-AAUP | 2621.49 | High |
| TCGA-HZ-8315 | 2629.69 | High |
| TCGA-F2-7276 | 2635.63 | High |
| TCGA-IB-7890 | 2640.61 | High |
| TCGA-F2-A44G | 2641.83 | High |
| TCGA-HZ-7925 | 2644.49 | High |
| TCGA-IB-8126 | 2672.59 | High |
| TCGA-IB-7886 | 2730.03 | High |
| TCGA-HZ-8003 | 2745.15 | High |
| TCGA-IB-7897 | 2807.37 | High |
| TCGA-IB-A5SO | 2901.59 | High |
| TCGA-Q3-A5QY | 2936.28 | High |
| TCGA-IB-AAUT | 2980.26 | High |
| TCGA-2L-AAQJ | 3068.81 | High |
| TCGA-FB-AAPQ | 3113.22 | High |
| TCGA-IB-7651 | 3241.11 | High |
| TCGA-IB-AAUS | 3274.87 | High |
| TCGA-HZ-7926 | 3279.51 | High |
| TCGA-IB-7893 | 3363.94 | High |
| TCGA-HZ-7918 | 3417.18 | High |
| TCGA-IB-7645 | 3510.19 | High |
| TCGA-IB-7647 | 3562.73 | High |
| TCGA-FB-AAPU | 3680.63 | High |
| TCGA-Z5-AAPL | 3958.4 | High |
| TCGA-F2-7273 | 4834.6 | High |
| TCGA-IB-7646 | 5423.35 | High |

**Table 1: Expression data from RNA and protein (RPPA) analyses for human pancreatic ductal adenocarcinoma patients in The Cancer Genome Atlas (TCGA) were extracted from cBioportal**. Expression of GNA13 RNA is categorized as high (GNA13 hi, upper 25%, n= 43) or low (GNA13 lo, lower 25%, n = 43). Table 1 lists patient IDs from TCGA used in our analysis.
