## Supplementary figures and images for "Gα13 loss promotes tumor progression in the KPC transgenic mouse model of advanced pancreatic cancer"

### SupplementalFigure1

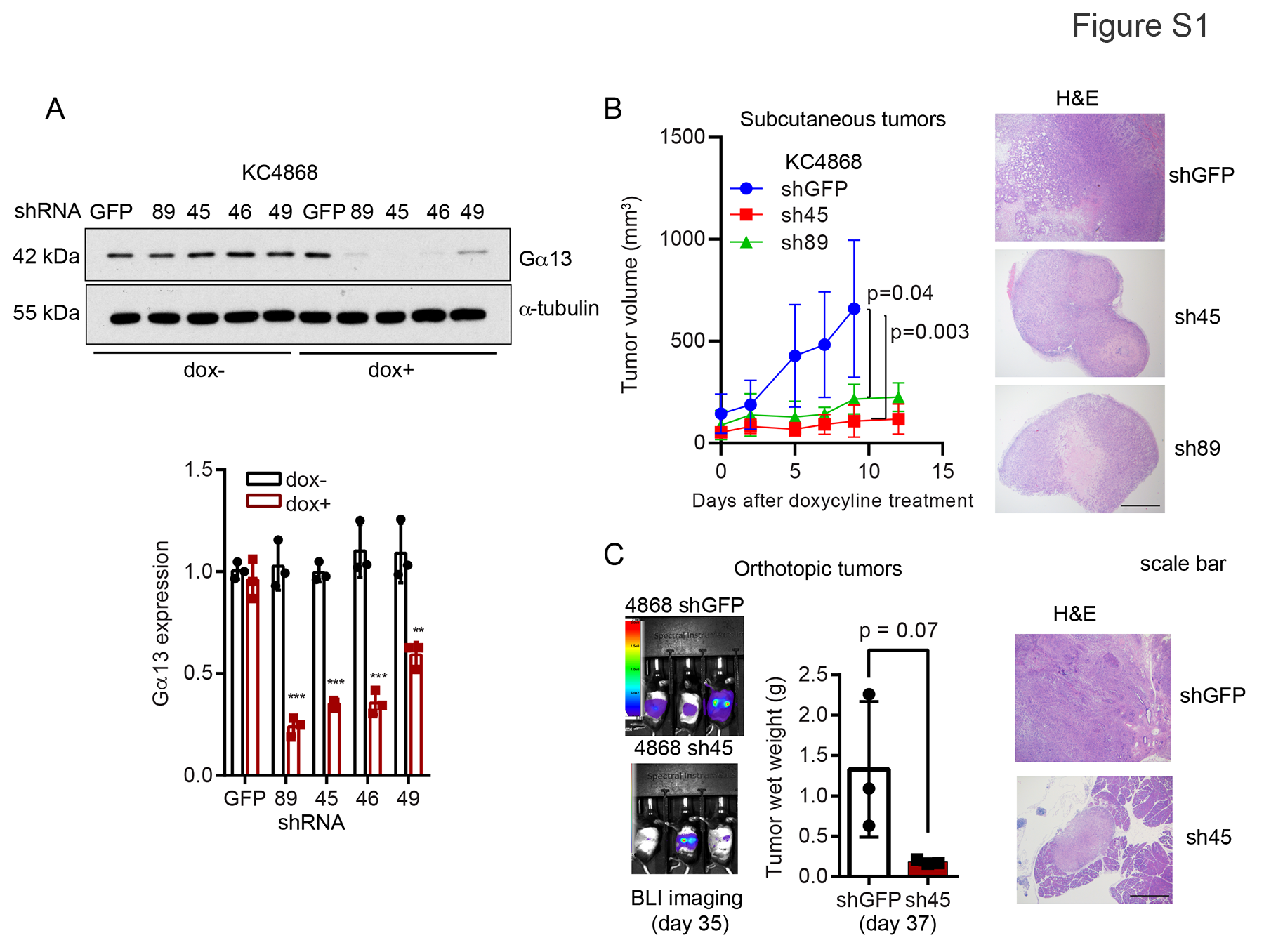
